# High accuracy DNA sequencing on a small, scalable platform via electrical detection of single base incorporations

**DOI:** 10.1101/604553

**Authors:** Hesaam Esfandyarpour, Kosar B. Parizi, Meysam R. Barmi, Hamid Rategh, Lisen Wang, Saurabh Paliwal, HamidReza Golnabi, Paul Kenney, Richard Reel, Frank Lee, Xavier Gomes, Seth Stern, Ashok Ramachandran, Subra Sankar, Solomon Doomson, Rick Ung, Maryam Jouzi, Ramya Akula Suresh Babu, Ali Nabi, Nestor Castillo, Raymond Lei, Mohammad Fallahi, Eric LoPrete, Austin Kemper, Srijeeta Bagchi, Robert Tarbox, Pallavi Choudhary, Hooman Nezamfar, Linda Hsie, Nicolas Monier, Tyson A. Clark, Eric Spence, Fei Yang, Benjamin Bronson, Gina Sutton, Caterina Schweidenback, John Lundy, An Ho, Narin Tangprasertchai, Anthony Thomas, Brian Baxter, Shankar Shastry, Anooshka Barua, Yongzhi Chen, Hamid Hashemzadeh, David Shtern, Eugene Kim, Christopher Thomas, Patrice Tanti, Ali Mazouchi, Erden Tumurbaatar, Jordan Nieboer, Christopher Knopf, Hien Tram, Vipal Sood, Sam Stingley, Megan Cahill, Sid Roy, Ky Sha, Bin Dong, Frank R. Witney, Ronald W. Davis

**Author notes:** corresponding author: Hesaam Esfandyarpour.

## Abstract

High throughput DNA sequencing technologies have undergone tremendous development over the past decade. Although optical detection-based sequencing has constituted the majority of data output, it requires a large capital investment and aggregation of samples to achieve optimal cost per sample. We have developed a novel electronic detection-based platform capable of accurately detecting single base incorporations. The GenapSys technology with its electronic detection modality allows the system to be compact, accessible, and affordable. We demonstrate the performance of the system by sequencing several different microbial genomes with varying GC content. The platform is capable of generating up to 2 Gb of high-quality nucleic acid sequence in a single run. We routinely generate sequence data that exceeds 99% raw accuracy with read lengths of up to 175 bp. Average quality scores remain above Q30 (99.9% raw sequencing accuracy) beyond 150 bp, with more than 85% of total bases at or above Q30. The utility of the platform is highlighted by targeted sequencing of the human genome. We show high concordance of SNP detection on the human NA12878 HapMap cell line with data generated on the Illumina sequencing platform. In addition, we sequenced a targeted panel of cancer-associated genes in a well characterized reference standard. With multiple library preparation approaches on this sample, we were able to identify low frequency mutations at expected allele frequencies.

## Introduction

The finished version of the human genome was released in 2004 (1), and soon after there was an initiative from the National Human Genome Research Institute (NHGRI) to sequence a human genome for $1000 (2). This initiative, in part, ushered in a period of development of a number of different DNA sequencing technologies (3-8). Today, a variety of commercial platforms exist that enable massively parallel, high throughput, and low-cost genomic sequencing, revolutionizing the nature of scientific and biomedical research. Despite continued technical improvements that have reduced the per base cost of DNA sequencing, the majority of sequencing is still carried out on large, expensive instruments with high costs per run (9).

We have developed a novel sequencing-by-synthesis approach that employs electrical detection of nucleotide incorporations. With electrical-based detectors, simple fluidics, no optics, and no robotics or moving parts, our automated sequencer is compact, robust, easy to use, and inexpensive. The instrument detects a steady-state signal, providing several key advantages over current commercially available sequencing platforms and allowing for highly accurate sequence detection. Sequencing is carried out on microfluidic chips that have a scalable number of detectors, supporting applications ranging from targeted sequencing of specific amplicons to genome-scale data collection. We believe that sequencing instruments that are compact, scalable, easy to use, and affordable to purchase and run will permit a more distributed model, in which genomic assays are put back into the hands of individual researchers. Here, we demonstrate the capabilities of the GenapSys sequencing platform in a variety of applications across multiple sample types.

## Results

### Detection of single base incorporation via electrical impedance modulation

An overview of the GenapSys sequencing platform is outlined in Figure 1. Briefly, a library of DNA fragments is clonally amplified via an isothermal amplification method onto primer-conjugated beads. Amplified beads are loaded onto the fluidic chamber on top of a fully integrated Complementary Metal-Oxide Semiconductor (CMOS) sequencing chip. The sequencing chip is then inserted into the GS111 instrument, which uses fluidics to inject reagents that will disperse evenly over the entire chip surface (Figure 2A). Nucleotides are injected one base at a time and incorporations are measured as electrical signals from each sensor that collect on the instrument’s Solid-State Device (SSD) storage unit. The incorporation data files are transferred in real time to a secure cloud-hosted server and storage environment. At the end of the run, the final FASTQ base call file becomes available.

**FIGURE 1:**
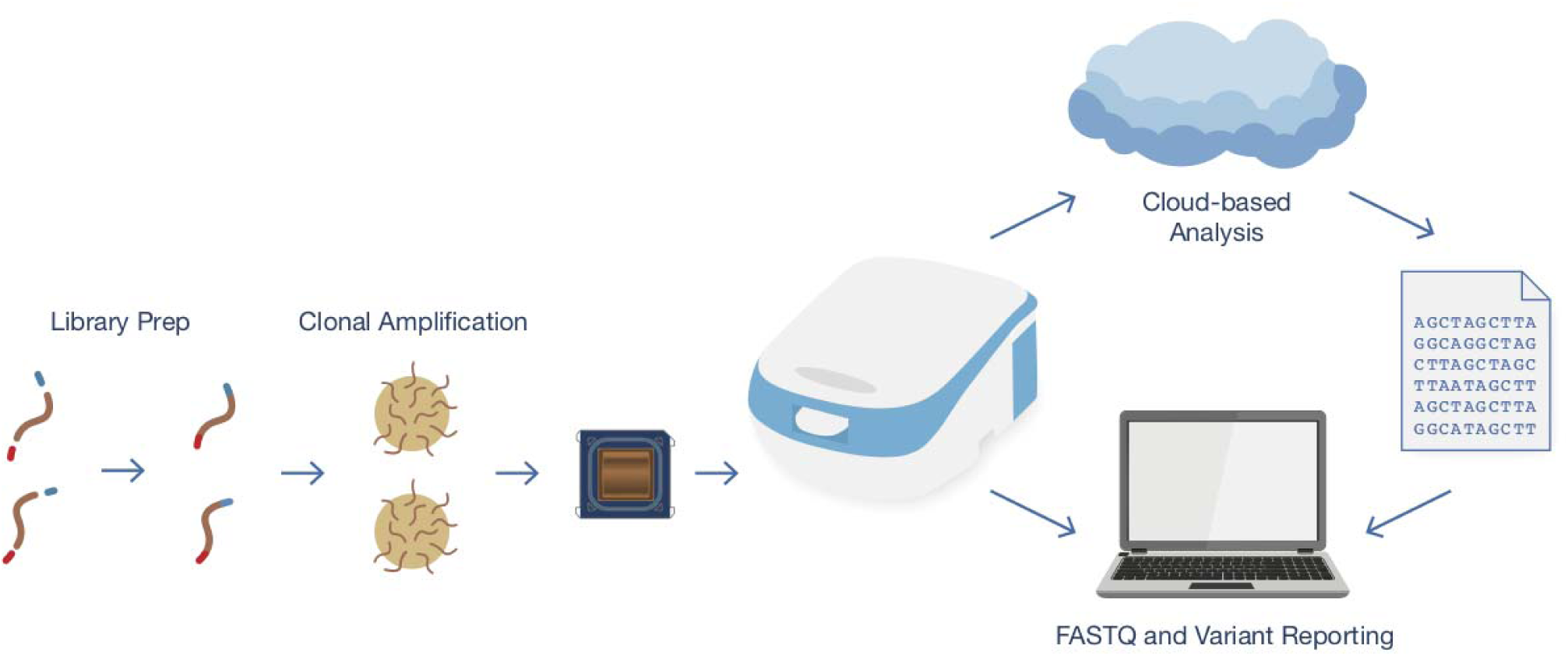
Overview of the Genapsys Sequencing Platform. An overview of the process for generating sequence data on the GenapSys sequencing platform. Libraries are clonally amplified onto beads, which are subsequently loaded onto the sequencing chip. Automated sequencing takes place on the GS111 instrument, which utilizes a cloud-based system for instrument control and data analysis, with the ultimate output being a FASTQ file of millions of individual DNA sequences with per-base quality scores.

**FIGURE 2:**
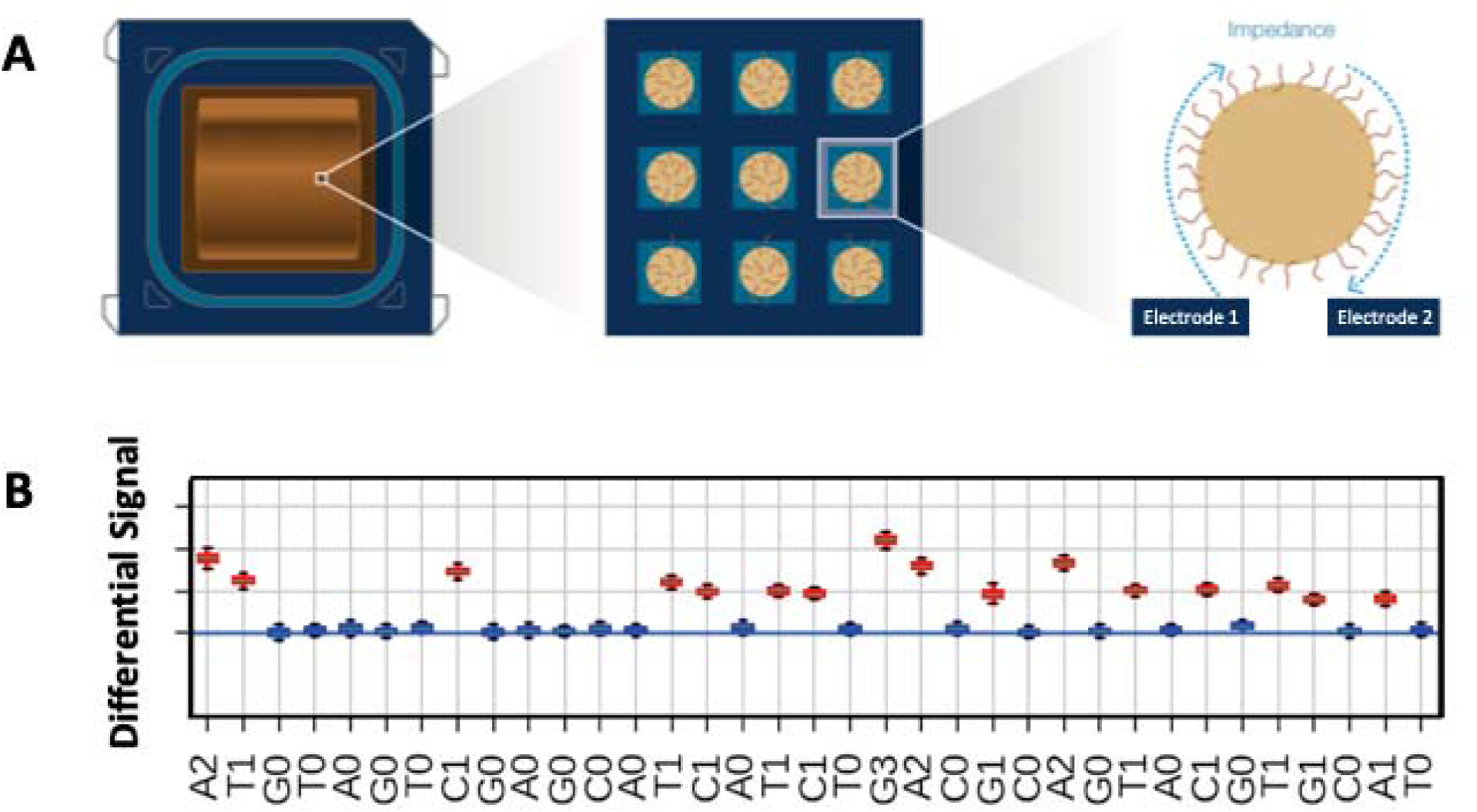
Sequencing Chip and Raw Data. (A) The CMOS sequencing chip contains millions of individual sensors, which are each loaded with a clonally amplified bead. The electrodes in each sensor are capable of measuring minute changes in impedance when nucleotides are incorporated opposite the bead-bound templates. (B) Raw data from each sensor is summarized as a differential signal for each nucleotide flow. This box plot represents the distribution of signal for a subset of nucleotide flows for a control template. Red boxes denote incorporating flows and blue boxes are flows with no nucleotide incorporation. The identity of the injected nucleotide and number of incorporated bases is shown below the plot.

Other Sequencing by Synthesis (SBS) approaches have been used successfully in several different commercial DNA sequencing platforms (6,7). The novelty of our approach lies in the detection modality. Other sequencing methods have relied upon transient signals such as pH or fluorescence-based detection. In contrast, our sequencer, measures a direct steady-state impedance-based electrical signal from clonally amplified DNA strands. The steady-state nature of our signal allows for multiple measurements over time to increase precision, which in turn improves the signal-to-noise ratio (SNR) and leads to higher accuracy base calls. The sequencing chip has a fully-integrated mixed-signal design, and uses a standard CMOS process.

The instrument is designed to be compatible with chips with varying number of sensors. To date, we have manufactured GenapSys sequencing chips with 1 million, 16 million, and 144 million sensors. All of the studies carried out in this work employ the mid-throughput chip with 16 million sensors. The instrument interfaces with the sequencing chip electronically and fluidically. The fluidics system of the instrument controls injection of different reagents including dNTP solutions, and buffers.

### Detection modality, signal processing and base calling

An example of the raw signal is shown in Supplementary Figure 1. Total signal of a sequencing bead increases with each successful incorporation (red line). A fraction of the sensors is purposefully loaded with amplified beads containing a sequencing primer blocked from amplification by virtue of a chemical modification at the primer’s 3’ end. Data from these sensors are used to normalize and remove any signal drift (blue line). The measured signal does not change significantly when a nucleotide is added that is not a match to the next DNA base in the template strand. When a nucleotide is incorporated, the measured impedance value of that sensor will jump creating a graph that resembles a staircase. Information on how the signal has changed upon injection of a given nucleotide can be summarized as a differential signal (Figure 2B). The magnitude of the differential signal correlates with the number of incorporated nucleotides. Thus, if the template contains more than one of the same consecutive base, the size of the signal change will be larger. Due to the steady-state nature of the impedance metric, the measured values do not significantly change over time. Thus, if desired, multiple measurements can be taken to improve precision. Figure 2B contains a representative example of acquired data showing the distribution of measured differential signal across multiple nucleotide flows for a single template sequence. Flows that are expected to have nucleotide incorporations are highlighted in red. Non-incorporating flows are shown in blue and do not deviate significantly from the baseline. The inferred sequence, including the number of nucleotides in each incorporation are shown below the plot (Figure 2B).

For each base, a quality score is predicted using a pre-trained Phred quality table (12). Low quality reads can be filtered based on the average quality score of their base calls. The substitution error rate is particularly low (∼ 0.01%) which compares favorably to other sequencing technologies (13-15). A breakout of the error types and a distribution of Q-score relative to position in the read for a typical run is shown in Supplementary Figure 2. Accuracy rates decrease slightly as read length increases, primarily due to phasing, similar to other technologies.

Homopolymer performance was evaluated based on data generated from a human whole exome-enriched library using Counterr (https://github.com/dayzerodx/counterr). This sample was used due to its richness of sequence contexts, providing more examples of homopolymer stretches of up to 10 consecutive nucleotides. Results from this analysis are visualized in Supplementary Figure 3. Using the human genome (hg38) sequence as a reference, we show our base calling results for each of the four bases for homopolymer lengths of 3 to 10. For comparison purposes, we show results from sequencing data from the same exome-enriched sample that had been sequenced using Illumina technology.

### Sequencing of Microbial Genomes

To test the performance of our platform, we used it to sequence the *E. coli* genome. Results are summarized in Table 1. The sequencing workflow is fully automated on the GS111 instrument with average run times that vary depending on the number of nucleotide flows. The average read length is 145 bp with base call lengths that range from ∼125 to 175 bp. The distribution of read lengths is shown in Supplementary Figure 4. Average coverage across the 4.69 Mb *E. coli* genome is 329x (Figure 3A-B), and the sequencing data shows relatively even coverage across the entire *E. coli* genome (Figure 3A). The average accuracy of the reads is 99.9% at 125 bp and remains well above 99% past 175 bp (Figure 3C). Nearly all of the bases have Q-scores greater than Q20 (99.0% accuracy) and more than 80% of the bases have Q-scores that exceed Q30 (99.9% accuracy).

**Table 1:** Sequencing Statistics (*E. coli*)

|  |  |
| --- | --- |
| Number of Reads | 11,253,515 |
| Number of Bases | 1,630,264,207 |
| Number of Mapped Reads | 10,586,584 |
| Number of Mapped Bases | 1,526,397,295 |
| Average Mapped Read Length | 145 bp |
| Raw Accuracy at Position 125 | 99.9% |
| Percentage of Bases with Q>20 | 99.9% |
| Percentage of Bases with Q>30 | 86.0% |
| Genome Size | 4.69 Mb |
| Average Coverage | 329x |

**FIGURE 3:**
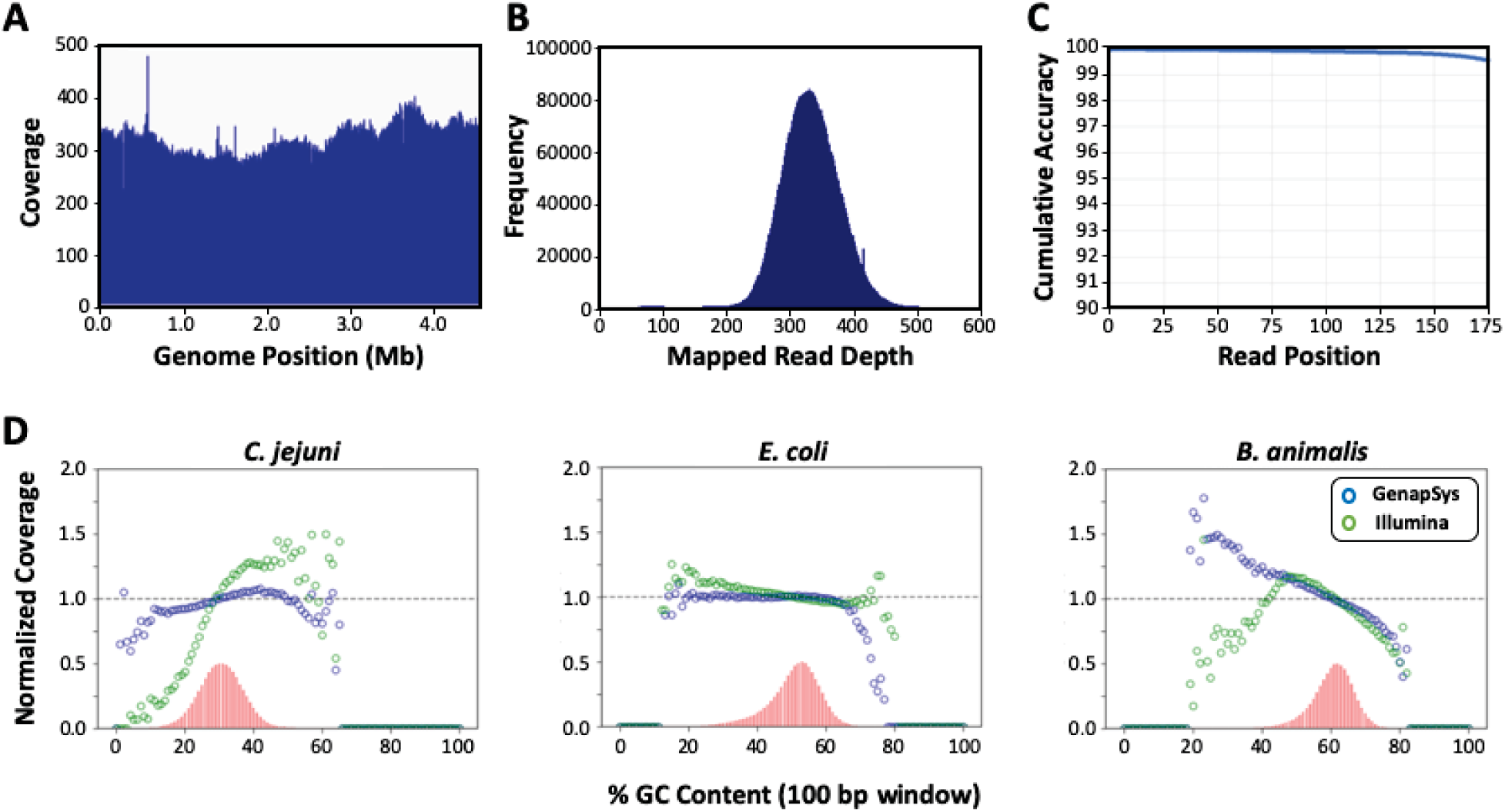
Microbial Genome Sequencing Statistics. (A) Coverage plot across the *E. coli* genome. (B) Histogram of mapped read depth for sequencing of the *E. coli* genome. (C) Average cumulative sequencing accuracy by read position. (D) Normalized coverage for sequencing of three different microbial genomes. Coverage is binned by percent GC content in a 100 bp window. The red histogram represents the abundance of each GC content bin in the genome. Blue circles are normalized coverage from the GenapSys platform and green circles are from data generated on the Illumina platform.

To further evaluate the system’s performance across a range of sequence contexts and a range of GC contents, we made libraries from additional microbial genomes. Details regarding the genomic DNA samples can be found in Supplementary Table 1. We selected microbial genomes with a range of average GC content. For example, *Campylobacter jejuni* is a low GC organism with an average GC content of about 30%. To address the higher GC content range, we sequenced *Bifidobacterium animalis subsp. Lactis* with an average GC content of 61%. Average coverage of each of these genomes exceeded 800x, with similar read lengths and accuracy statistics to the *E. coli* run. Statistics from the sequencing runs are outlined in Supplementary Figure 5. Figure 3D shows the distribution of sequencing coverage broken down into bins of GC content. For comparison, we also show Illumina data generated from the same genomic DNA. This data shows that the sequencing system is robust and relatively uniform across a wide variety of sequence contexts and GC content. While there is some decrease in normalized coverage at the GC content extremes, the bias is consistent with what is typically observed for sequencing technologies that rely upon amplification (16).

### SNP Discovery in a Human Exome Sample

As a demonstration of the technology applied to human samples, we carried out whole exome sequencing of DNA from the extremely well characterized human cell line NA12878. NA12878 is a lymphoblastoid cell line (LCL) prepared from a normal female Caucasian participant of the international HapMap project (17). This DNA sample has served as human reference standard for the Genome in a Bottle (GIAB) Consortium and has been well characterized using a variety of different genomic technologies (18).

Targeted sequencing libraries made from NA12878 genomic DNA were generated via probe-based capture using the IDT xGEN Exome Research Panel (v1.0). The exome library was sequenced to approximately 115x average coverage across the 39 Mb targeted region. Approximately 81% of the sequencing reads mapped to the target region with relatively uniform coverage (Supplementary Table 2). The read length and accuracy are similar to the *E. coli* run detailed above.

**Table 2:** Low Frequency Variant Detection via Hybrid Capture Probe-based Targeted Sequencing

| Gene | Variant | Expected | GenapSys | Illumina |
| --- | --- | --- | --- | --- |
| EGFR | T790M | 1.0 | 1.0 | 1.0 |
| EGFR | delE746-A750 | 2.0 | 2.5 | 1.7 |
| EGFR | L858R | 3.0 | 3.0 | 2.8 |
| KRAS | G12D | 6.0 | 3.5 | 4.3 |
| PIK3CA | E545K | 9.0 | 3.8 | 6.6 |
| KIT | D816V | 10.0 | 8.5 | 6.9 |
| BRAF | V600E | 10.5 | 7.8 | 8.9 |
| NRAS | Q61K | 12.5 | 10.4 | 10.6 |
| KRAS | G13D | 15.0 | 14.1 | 15.4 |
| PIK3CA | H1047R | 17.5 | 14.2 | 15.2 |
| EGFR | G719S | 24.5 | 23.2 | 23.3 |

We identified single nucleotide variants (SNPs) relative to the human genome reference (hg38), using DeepVariant (20). DeepVariant has shown better performance than conventional tools and is, reportedly, more generalizable to the error profiles of different sequencing technologies (20). When carrying out the variant analysis with a focus on the high confidence regions from the GIAB consortium (v3.3.2), the GenapSys sequencing data of the NA12878 exome generated 21,013 SNP variants. To benchmark the performance of the GenapSys platform for SNP detection, we subjected the same library to Illumina sequencing with an average coverage of approximately 526x. Using the pre-trained DeepVariant model from Google (v0.7.2), the Illumina data identified 21,108 SNPs. The overlap between the two platforms was extremely high. This is illustrated in Figure 4A, where the Venn diagram shows sensitivity of 99.3%, precision of 99.8% and an F1 score of 99.5%, with low levels of both false positives and false negatives for the GenapSys data relative to the Illumina data. If we consider the GIAB high confidence call set as ground truth, we maintain high concordance with 99.2% sensitivity, 99.9% precision, and an F1 score of 99.5% as shown in Figure 4B. In Supplementary Figure 6 we show the results for the test regions of the DeepVariant analysis. The ratio of transition to transversion SNPs (TiTv ratio) is 2.98. Overall, this compares very favorably to similar comparisons done between different sequencing platforms (19).

**FIGURE 4:**
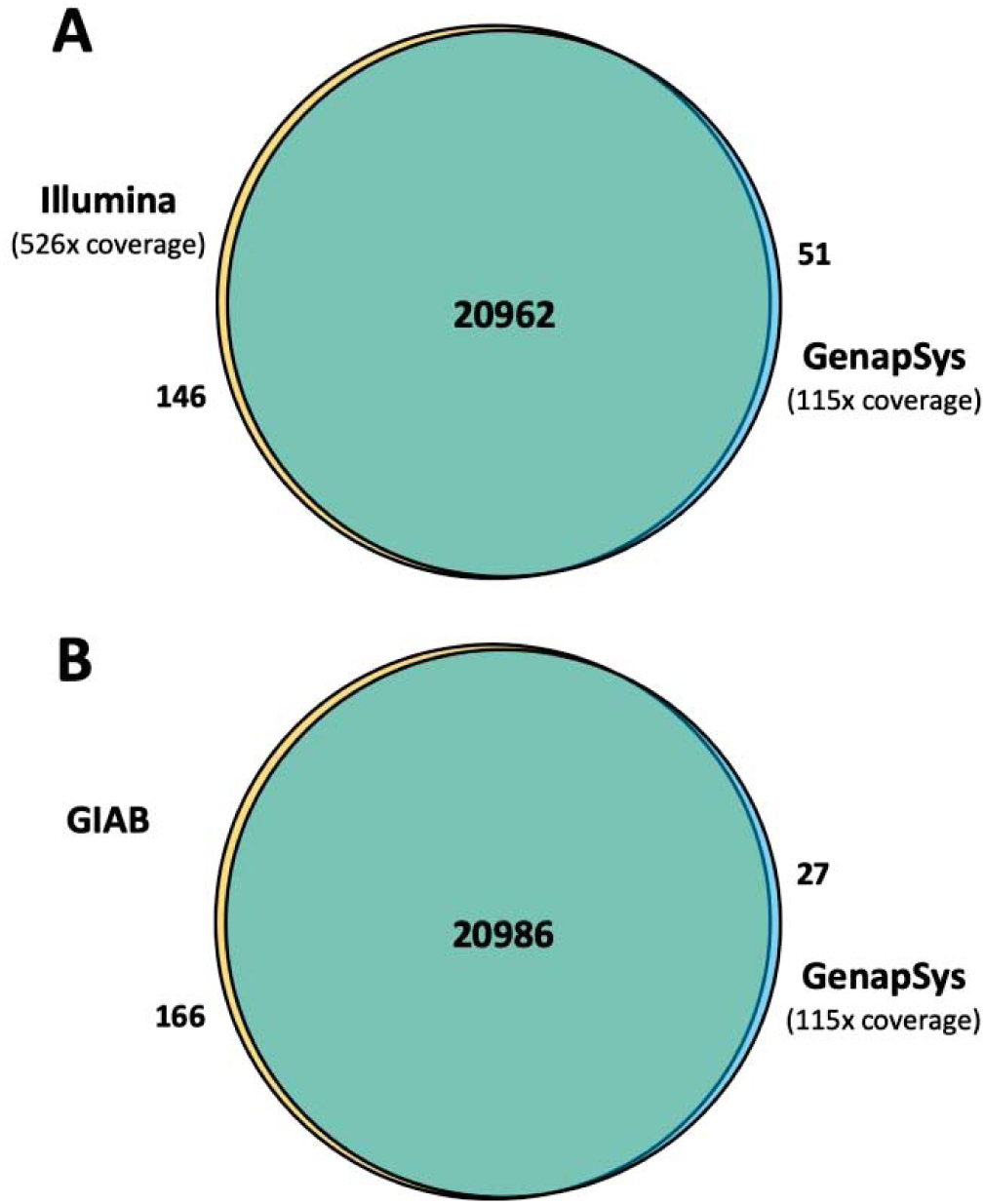
Human Exome Variant Concordance. Venn diagrams illustrating the overlap of SNPs identified in the NA12878 Exome sample from GenapSys data. DeepVariant was used to call variants. (A) GenapSys SNPs relative to SNPs called from the Illumina dataset using high confidence regions from the GIAB consortium. (B) GenapSys SNPs relative to the GIAB high confidence call set.

We also evaluated other variant calling methods. When using standard analysis methods such as BCFtools in conjunction with the same high confidence regions we achieve similar concordance. The results are shown in Supplementary Figure 7. The Venn diagrams show significant overlap between variant calls from the GenapSys data and libraries sequenced with Illumina and the GIAB high confidence variants. However, as a more generalizable variant calling algorithm, the DeepVariant approach could be preferable for use with GenapSys sequencing data over other commonly used tools such as GATK (21), which is specifically tuned to the particular error modes present in Illumina sequencing data.

### Low Frequency Variant Detection in a Targeted Human Cancer Reference Sample

Quantitation of low frequency somatic mutations in cancer samples requires a sequencing platform with high accuracy. To evaluate the performance of the GenapSys platform for this application, we performed targeted sequencing of a quantitative multiplex reference standard (Horizon Discovery HD701) using a focused cancer probe panel from IDT (xGen Pan-Cancer v1.5). The genomic DNA sample is a mixture of genomic DNA from 3 cell lines, HCT116, RKO and SW48, with verified oncology-relevant mutations, with allele frequencies that range from 1% to 24.5%. One sequencing run generates more than sufficient coverage for detection of low frequency alleles. The on-target rate across the 800 Mb panel was more than 64% and read length and accuracy statistics were similar to the *E. coli* runs (Supplementary Table 2).

Table 2 shows the concordance of our SNP variant calls with the expected allelic frequencies for 11 different mutations. The allele frequencies measured by the GenapSys sequencing platform closely match the expected allele frequencies of the quantitative multiplex reference standard, determined by the vendor using Droplet Digital PCR. Mutant alleles with frequencies as low as 1.0% were correctly identified. As an example, aligned reads visualized in the IGV browser for EGFR variant L858R are shown in Supplementary Figure 8. There are 925 reads that map to this genomic location and 28 of them show the variant, correlating well to the expected allele frequency of 3.0%. There is a relatively balanced proportion of forward and reverse reads from both the mutant and wild type alleles. As a benchmark, we sent the same library out for sequencing on the Illumina sequencing platform and used the same analysis methods on both sets of sequencing data. The measured allele frequencies are similar. A similar analysis was carried out on cancer gene-enriched libraries made from the OncoSpan quantitative reference DNA (Horizon Discovery). A scatter plot of allele-frequencies from GenapSys sequencing data relative to Illumina data demonstrating high correlation (R^2^ = 0.98) is shown in Supplementary Figure 9. Taken together, these data demonstrate comparable performance of the GenapSys sequencing platform with well-established methods, for applications requiring low frequency variant detection.

We also investigated low frequency mutation detection with a library preparation approach using multiplex PCR amplicons. The Ion AmpliSeq Hotspot Cancer Panel v2 consists of 207 primer pairs and generates amplicons targeting genomic ‘hot spot’ regions that are commonly mutated in human cancers. An amplicon library was generated using the AmpliSeq panel with the Horizon HD701 gDNA sample. Sequencing of this library using the GenapSys platform led to the identification of the expected mutations, including low-frequency mutations. Figure 5 shows the correlation of expected allele frequency to that measured by the GenapSys sequencing platform. The correlation is very high with an R^2^ of greater than 0.99.

**FIGURE 5:**
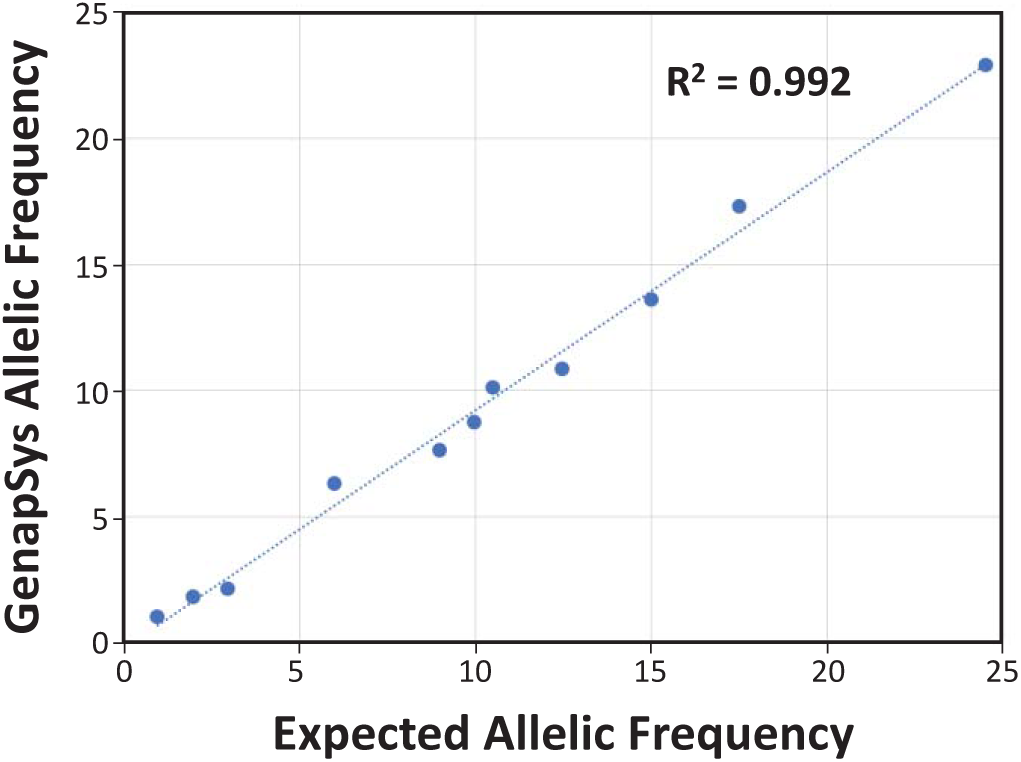
Low Frequency Variant Correlation via Amplicon-based Targeted Sequencing. Scatter plot showing high correlation between measured and expected allelic frequency for 11 variants from the HD701 Quantitative Multiplex Reference Standard. The x-axis represents the expected allelic frequency in the HD701 gDNA as measured by ddPCR and the y-axis represents the measured allelic frequency for AmpliSeq Cancer Hotspot Panel v2 targeted sequencing on the GenapSys platform. The Pearson correlation coefficient is 0.992.

## Conclusions

We have demonstrated that the GenapSys sequencing platform is capable of generating very accurate DNA sequence data, enabling its use for a range of applications such as whole genome sequencing, exome sequencing and low frequency variant detection. It routinely generates sequence data that exceeds >80% of bases >Q30 with average read lengths of approximately 150 bp. The system’s architecture, including CMOS-based electronic detection, an absence of moving parts and optics, minimal computational requirements, and simple fluidic controls, allows for an instrument that is compact, scalable, and affordable.

As with many other sequencing technologies, we believe this platform is capable of rapid technological improvements that will increase accuracy, throughput, and speed. In particular, the CMOS technology can make it feasible to have significantly higher sensor densities, and hence a big increase in throughput. The instrument’s flexible architecture accommodates such improvements by modifying the consumables without substantially altering the instrument itself. Therefore, additional sequencing throughput can be achieved without significant instrument modifications or run time increases, thereby dramatically improving the cost per base.

Further improvements to the sequencer and workflow have the potential to improve the performance and usability of the platform. Enhanced nucleotide modification chemistries can increase the signal to noise ratio, improve accuracy and allow for longer read lengths. Better optimization of flow times could reduce the duration of sequencing runs. Automation and integration of sample preparation prior to sequencing could further increase the ease of use. Our detection architecture is compatible with techniques that do not require beads, in which the sequencing libraries are clonally amplified directly on the surface of the chip. These changes will allow all steps of the sequencing process to be automated in a single instrument.

In the past two decades, a few DNA sequencing technologies have been developed and made commercially available. Currently, the vast majority of DNA sequence data is generated using optical based approaches. Dramatic improvements in throughput have made it possible, for example, to re-sequence a human genome at a cost approaching $1000 per sample. However, these sequencing runs are carried out on large, expensive instrumentation with high per run costs. Though a reduction in cost per base has been enabled by extremely high output runs and multiplexing of multiple samples into a single run, the instrument price and run price is still out of reach for many individual researchers.

One of the key innovation of GenapSys’ platform is the steady-state electrical detection of base incorporations. Combining this with fully integrated CMOS chips, enables the system to be small, accessible, and affordable. Its flexible architecture allows the system to be highly scalable, with potential chip throughputs ranging from one million to hundreds of millions of sensors, all compatible with an instrument with a small footprint and ease of use. The detection modality can also be extended to other types of nucleic acids, proteins, and live single cells using the same platform. We believe our sequencing technology will ultimately extend the reach of genomics to a wider audience and a wider range of applications.

## Materials and Methods

### Library Preparation

Genomic DNA samples were purchased from various commercial sources. A table of genomic DNA samples used in this study can be found in Supplementary Table 1. Sequencing libraries were generated by random shearing of genomic DNA followed by ligation of custom adapters using the Kapa HyperPrep Kit (Roche). Briefly, approximately 1 μg of genomic DNA was acoustically sheared to ∼200 bp using a M220 Focused-ultrasonicator (Covaris). Sheared DNA was end repaired, A-tailed, and ligated to custom adapters (Integrated DNA Technologies) using the Kapa HyperPrep Kit per manufacturer’s recommendations. Ligated fragments were purified with AMPure XP beads (Beckman Coulter) and size-selected on a Pippin Prep (Sage Science). Size-selected libraries were PCR amplified using the Kapa HiFi PCR Kit (Roche) and purified using AMPure XP beads. The qualities of final libraries were assayed using a Bioanalyzer High Sensitivity DNA Kit (Agilent).

### Target Capture and Multiplex PCR Amplicon Panels

Targeted sequencing libraries were enriched from standard sequencing libraries using xGen Lockdown Panels (Integrated DNA Technologies). Genomic libraries from the NA12878 cell line (Coriell) were enriched with the xGen Exome Research Panel v1.0, and libraries from the Horizon Quantitative Multiplex Reference Standard gDNA (HD701) and OncoSpan (HD827) were enriched with the xGen Pan-Cancer Panel v1.5. Enrichment was carried out per manufacturer’s recommendations using the xGen Hybridization and Wash Kit (Integrated DNA Technologies) and custom xGen Blocking Oligos complementary to our custom adapter sequences. Enriched libraries were subsequently amplified using the Kapa HiFi PCR Kit and purified using AMPure XP beads.

Multiplex PCR Amplicon panels for Cancer were generated using the Ion AmpliSeq Cancer Hotspot Panel v2 (ThermoFisher Scientific). Multiplex PCR was performed on 10 ng of genomic DNA from the HD701 reference standard, followed by partial digestion of primer sequences by the FuPa reagent, according to the manufacturer’s recommendations. The amplified products were purified using AMPure XP beads, end-repaired, A-tailed, and ligated to custom adapters as above. Ligated fragments were amplified by PCR and purified using AMPure XP beads, to give rise to the final library.

### Clonal Amplification and DNA Sequencing

Library molecules were isothermally clonally amplified onto primer conjugated beads using manufacturer’s recommendations (GenapSys).

Clonally amplified and enriched beads were mixed with reference beads prior to addition to the sequencing chip. Reference beads are amplified beads that have been annealed with a 3’ blocked sequencing primer. Beads were loaded onto the sequencing chip which has an array of sensors, each capable of capturing one bead. GenapSys Sequencing polymerase was injected into the chip and incubated for 5 minutes to allow for binding to the DNA templates per manufacturer’s recommendations.

Sequencing begins with electrical calibration of the chip, followed by serial injection of individual nucleotide buffers. For each injection, the chip was incubated for several seconds to allow for nucleotide incorporation, then the electrical signal of individual sensors was captured. The chip was washed, and this process was then repeated using a different nucleotide. To sequence of at least 150 base pairs, approximately 250-300 nucleotide flows were required.

### Signal Processing and Base Calling

Sequence data was generated using version 2.9.6 of the GenapSys analysis pipeline. The initial step of data analysis was sensor identification where sensors with beads are differentiated from those without a bead based on initial sensor calibration (Supplementary Figure 10A). The population of sensors with bead were further grouped into sequencing and reference based on the characteristics of their measured signal over a predetermined number of nucleotide flows (Supplementary Figure 10B). Reference sensors are those with blocked primer beads and were used to subtract the background noise from the sequencing sensors’ signal.

Base calling was carried out using the GenapSys proprietary base calling pipeline. Data from each sequencing sensor for each nucleotide flow was evaluated relative to the reference sensors. The base caller identified the number of incorporations in the case of homopolymer sequence stretches. Detailed analysis of the homopolymer calling performance is shown in Supplementary Figure 3.

### Sequence Alignment and Variant Detection

The reference genomes were indexed using BWA (v0.7.17) and BWA-MEM was used to align the single-end reads to the reference genome (22). Sequencing reads were aligned to the hg38 reference genome using BWA-MEM. To call variants, DeepVariant (20) and BCFtools were used. Variant analysis was carried out using the DeepVariant (v0.8.0) approach. The pre-trained DeepVariant Illumina WES model from Google was further trained on the GenapSys NA12878 exome dataset with 60% training, 20% cross-validation, and 20% test. Additionally, the ‘mpileup’ command in BCFtools (v1.9) was used to generate the genotype likelihoods at each genomic position with sufficient coverage. The BCFtools ‘call’ command was then used to call the variant sites using default settings. The BCFtools ‘isec’ command was used for variant evaluation.

### Data Availability

Sequence data has been deposited into the Sequence Read Archive (SRA) under BioProject accession number PRJNA529876.

## Supplementary Material

Supplementary Figure 1: Steady-State Signal of Individual Base Incorporations

Supplementary Figure 2: Error Types and Q-Score Distribution

Supplementary Figure 3: Homopolymer Performance

Supplementary Figure 4: Base Call Length Distribution

Supplementary Figure 5: Microbial Genomes

Supplementary Figure 6: DeepVariant SNP Concordance of Test Regions

Supplementary Figure 7: BCFtools Analysis of SNPs in NA12878 Exome Data

Supplementary Figure 8: Low Frequency Variant Read Visualization

Supplementary Figure 9: Low Frequency Variant Detection via Hybrid Capture Probe-based Targeted Sequencing

Supplementary Figure 10: Sensor Identification and Active Sensor Determination

Supplementary Table 1: Genomic DNA Samples

Supplementary Table 2: Targeted Sequencing Statistics

## Acknowledgements

We would like to thank current and past members of the GenapSys team who contributed to the development of the sequencing technology.

## Author Contributions

H.E. invented the technology and directed the project. T.A.C., S.P., A.N., M.F. and H.E. designed the studies and T.A.C. wrote the manuscript with input and support from other co-authors. K.B.P., H.R., A.R., N.M. and H.T. designed the CMOS sequencing chip. L.W., H.R.G., R.R., F.L., S.S., N.C., R.L., E.S., E.K., J.N. and C.K. developed the instrument. S.P., X.G., A.K., L.H., F.Y., C.S., J.L., A.B., C.T., E.T. and K.S. developed wet lab methods. M.R.B., M.J., G.S., N.T., and A.T. performed the sequencing experiments. P.K. and B.B. developed surface chemistry methods. M.R.B, K.P.B, M.J., S.S., R.A.S.B., P.C., and S.S. developed the sequencing chemistry. R.U., B.B., A.H., Y.C., H.H., D.S., and A.M. developed the software. A.N., M.F., E.L, S.B, H.N., and B.D. developed algorithms and analyzed the data. S.D., R.T., P.T., V.S., S.S., M.C., S.R. and F.R.W. provided support to the R&D team.

## Author Information

The authors declare competing financial interests: all authors are current or former employees or contractors that work for GenapSys, Inc. R.W.D. is a member of GenapSys’s Scientific Advisory Board.

