## Supplemental Material for "High accuracy DNA sequencing on a small, scalable platform via electrical detection of single base incorporations"

**SUPPLEMENTARY FIGURE 1:** Steady-State Signal of Individual Base Incorporations

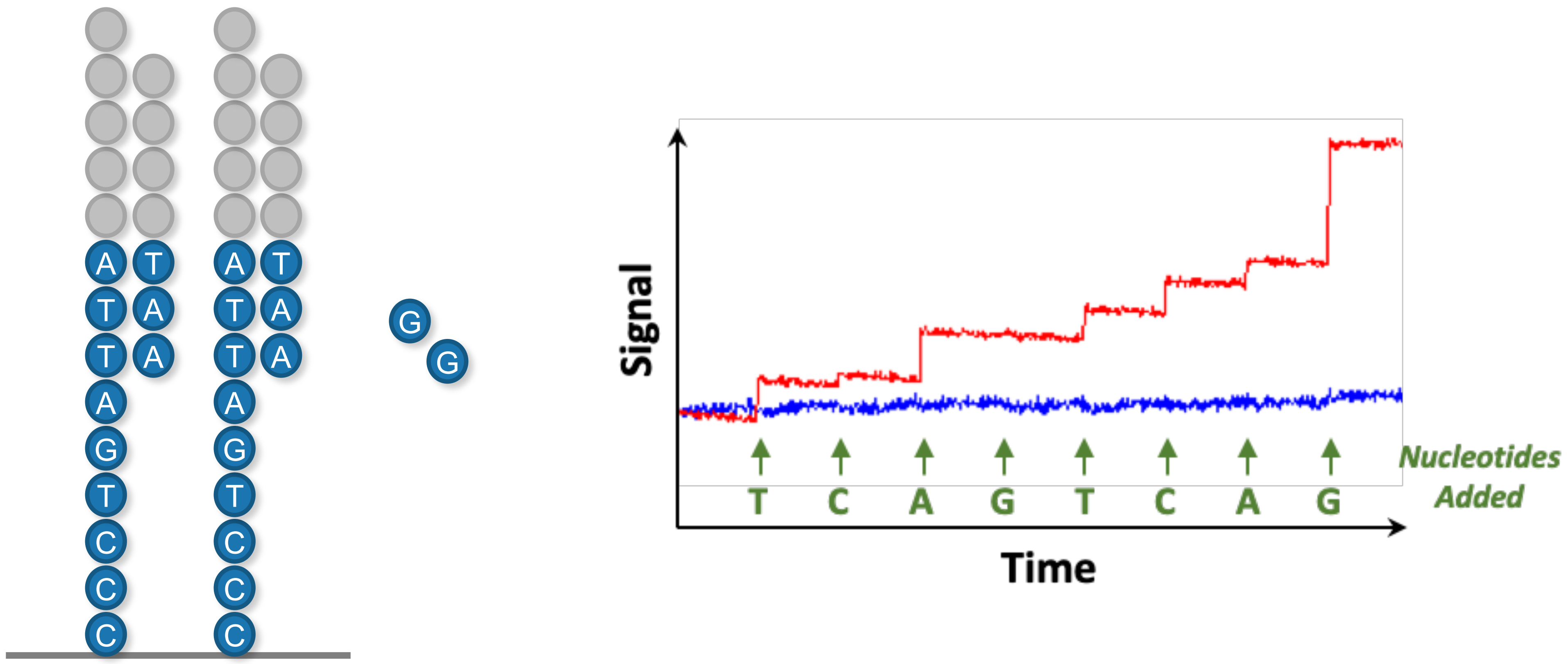

**SUPPLEMENTARY FIGURE 2:** Error Types and Q-Score Distribution

***E. coli* Read Accuracy* = 99.9%**

| Substitution Errors | Deletion Errors | Insertion Errors |
|---|---|---|
| **0.01%** | **0.09%** | **0.03%** |

** at base position 100*

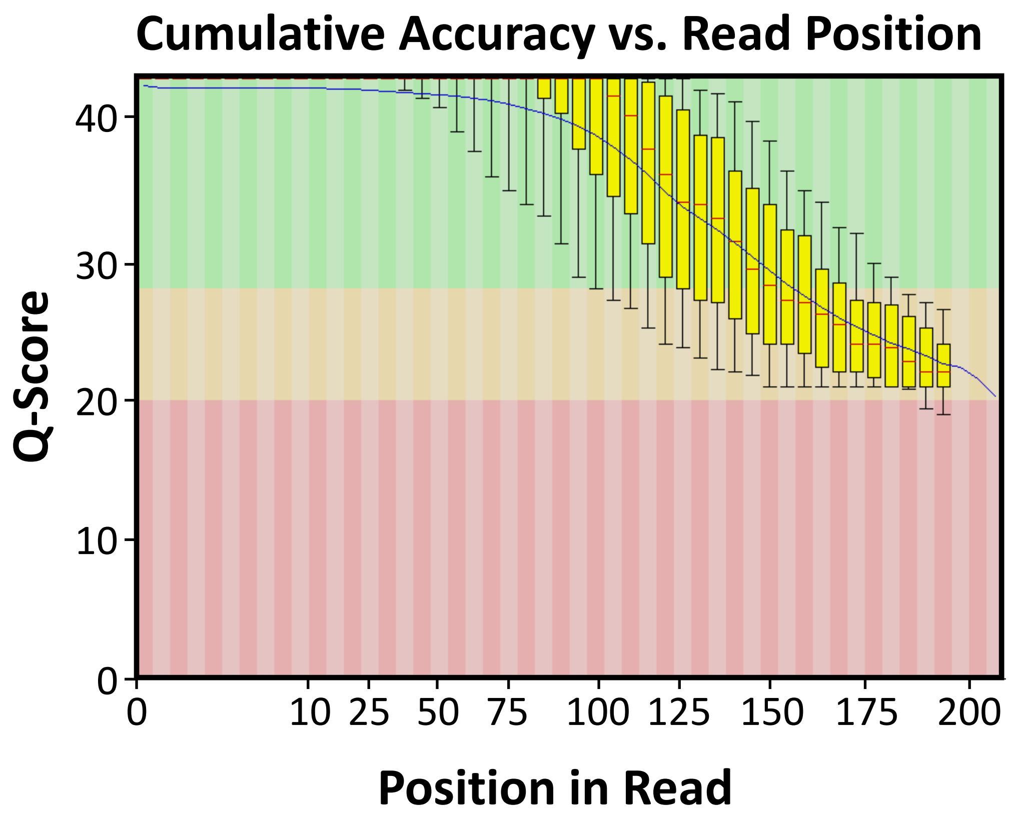

### **SUPPLEMENTARY FIGURE 3:** Homopolymer Performance

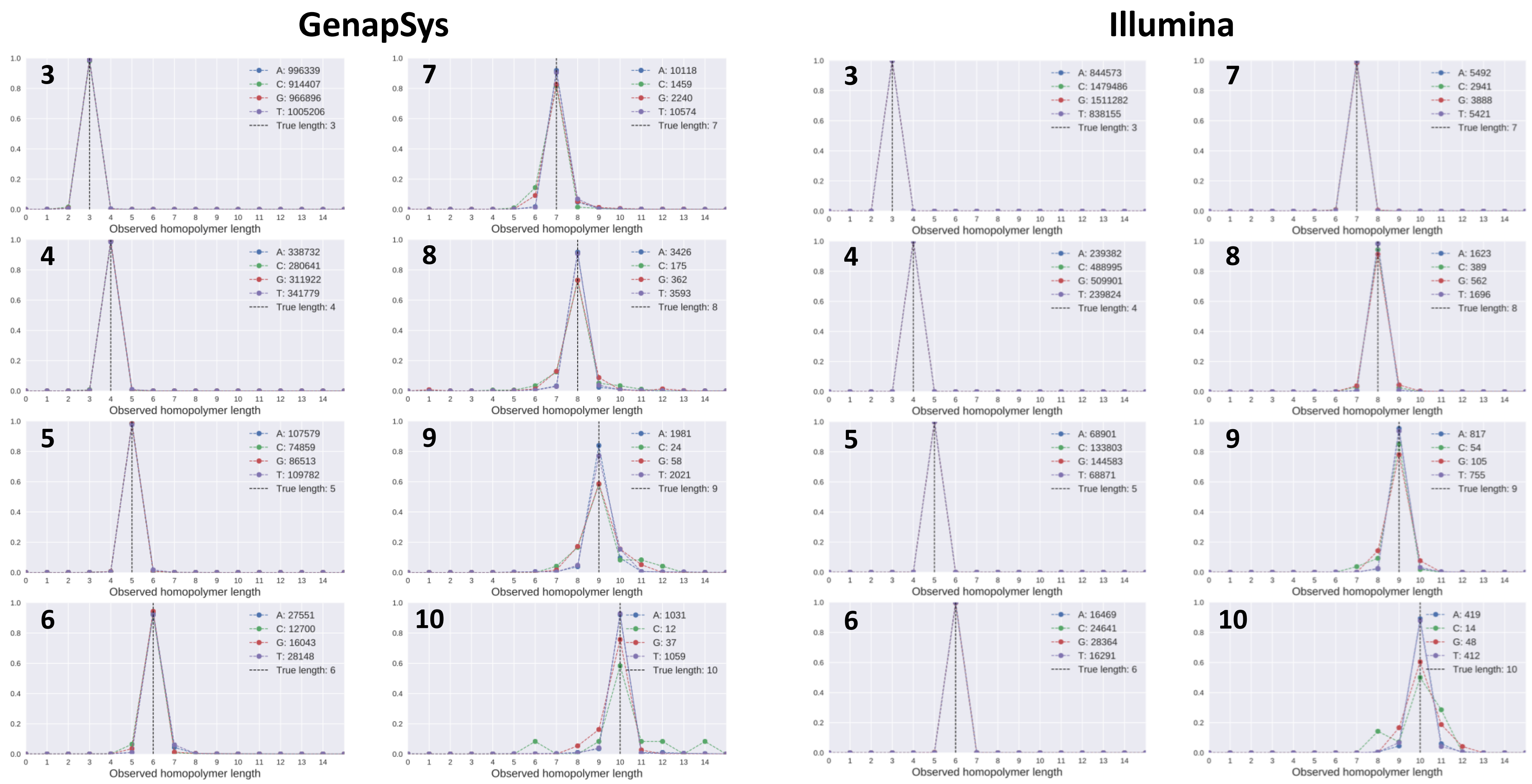

**SUPPLEMENTARY FIGURE 4:** Base Call Length Distribution

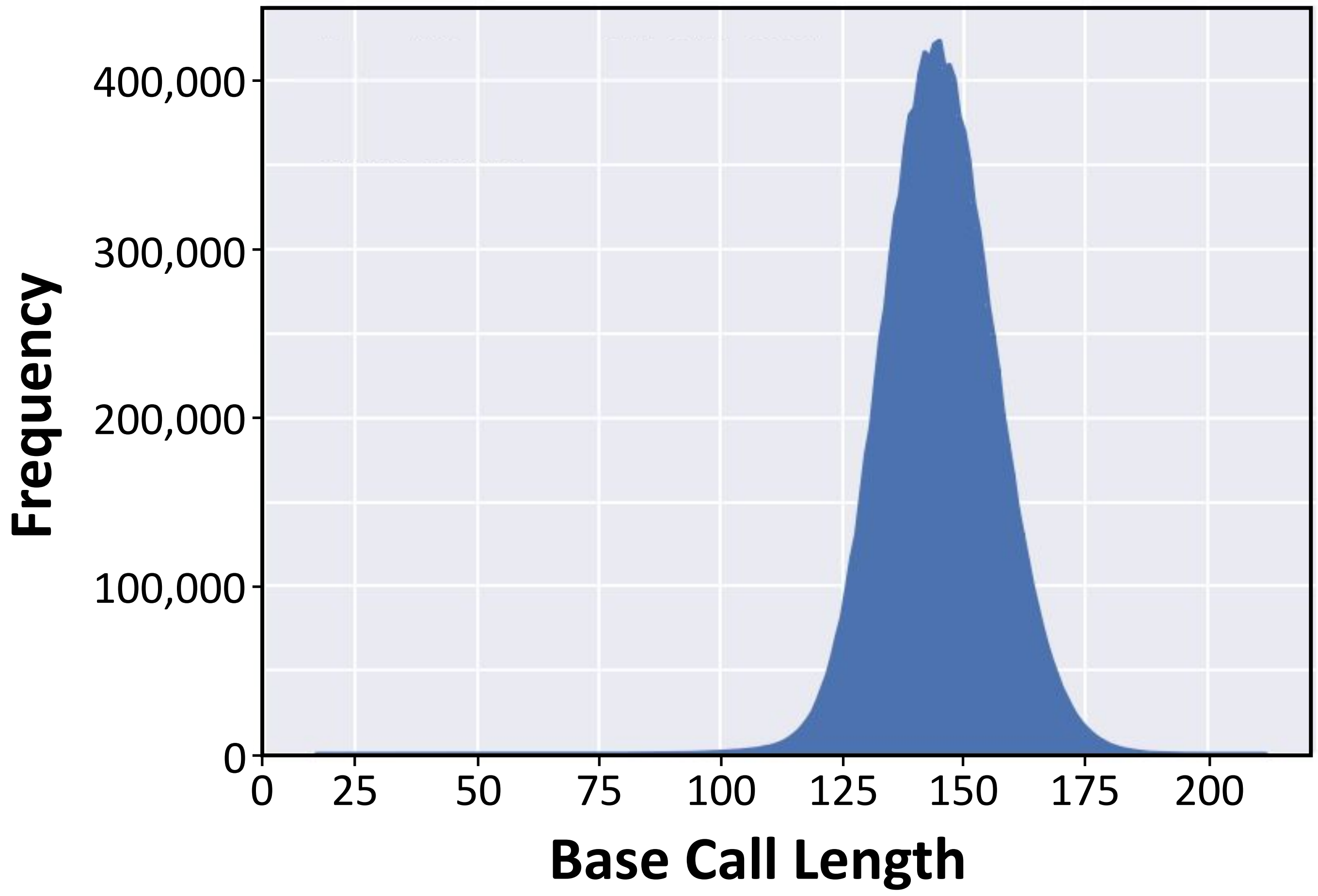

**SUPPLEMENTARY FIGURE 5:** Microbial Genomes

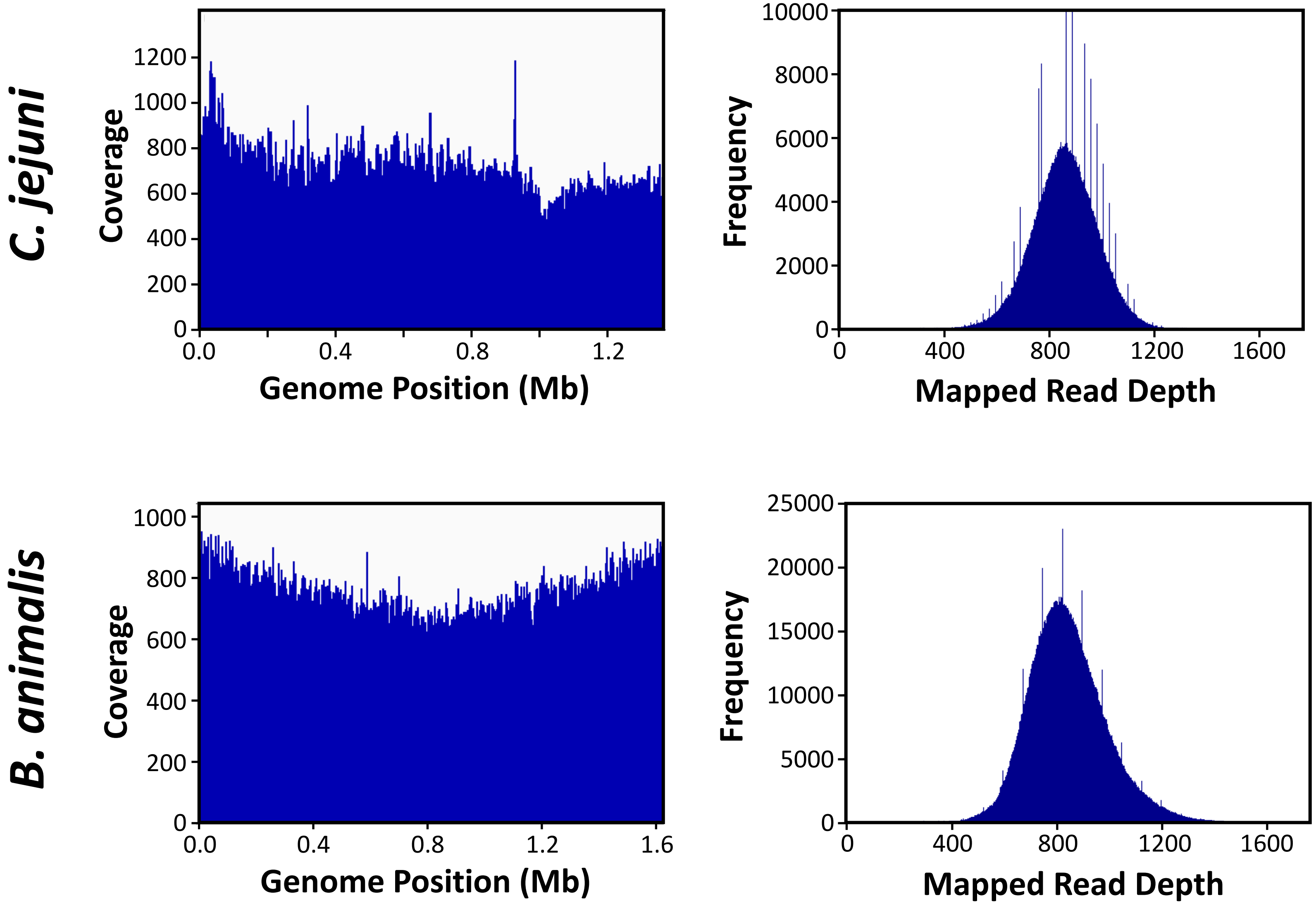

**SUPPLEMENTARY FIGURE 6:** DeepVariant SNP Concordance of Test Regions

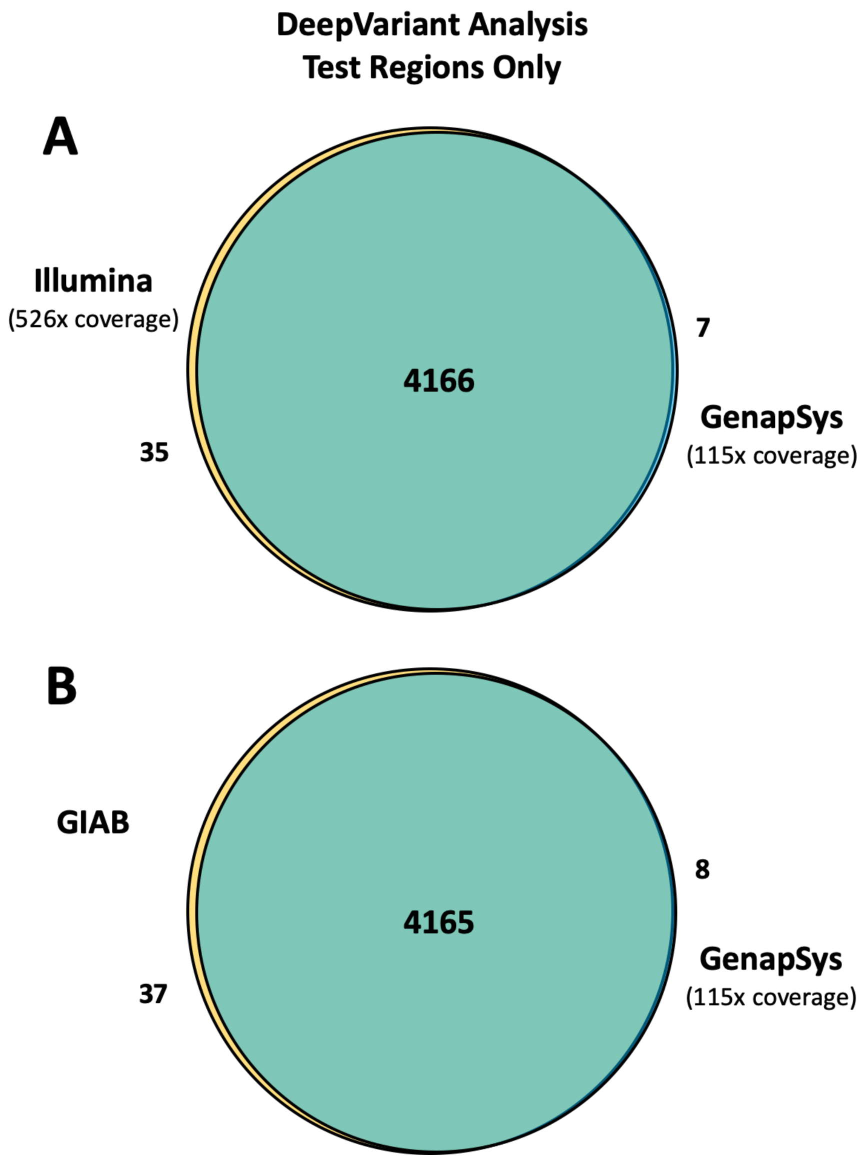

**SUPPLEMENTARY FIGURE 7:** BCFtools Analysis of SNPs in NA12878 Exome Data

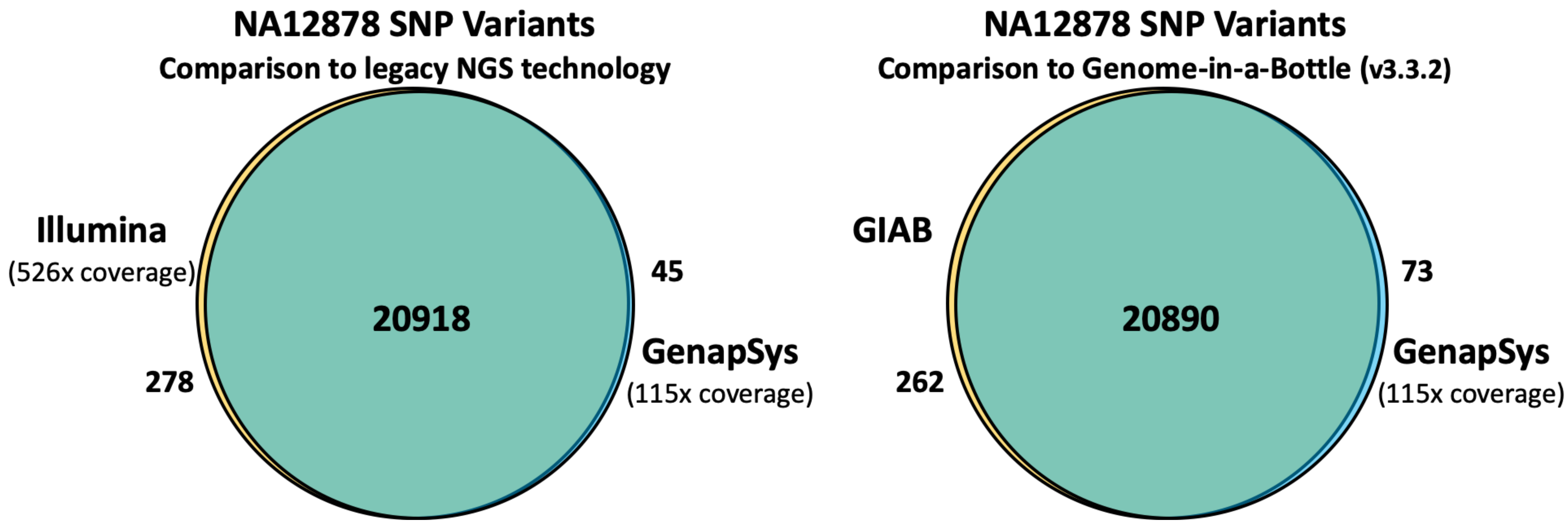


| Metric | GenapSys vs Illumina | Genapsys vs GIAB |
|---|---|---|
| Sensitivity | 98.7% | 98.8% |
| Precision | 99.8% | 99.7% |
| F1 Score | 99.2% | 99.2% |

**SUPPLEMENTARY FIGURE 8:** Low Frequency Variant Read Visualization

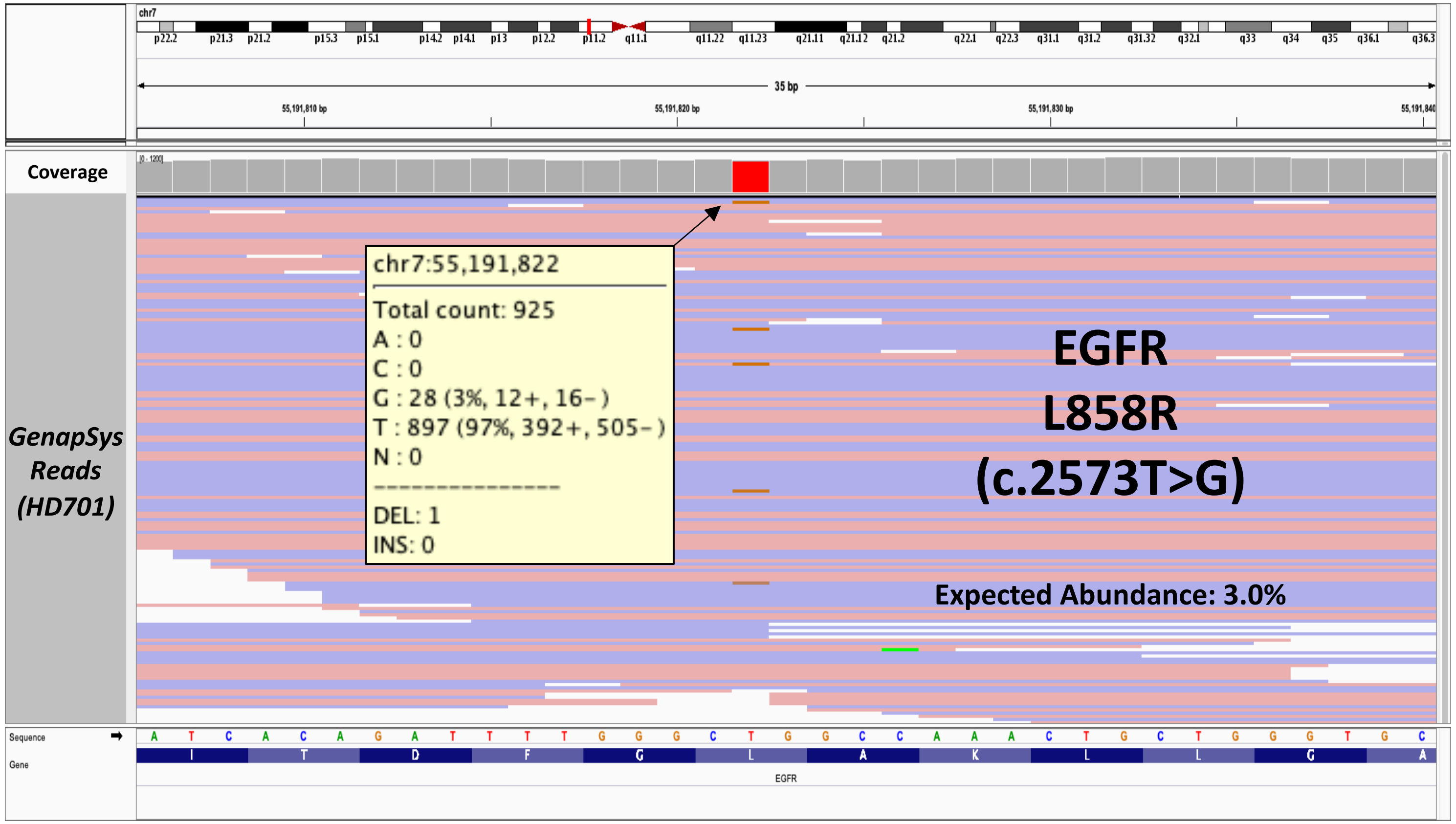

**SUPPLEMENTARY FIGURE 9:** Low Frequency Variant Detection via Hybrid Capture Probe-based Targeted Sequencing

OncoSpan (HD827) HotSpots in IDT xGen Pan Cancer Panel

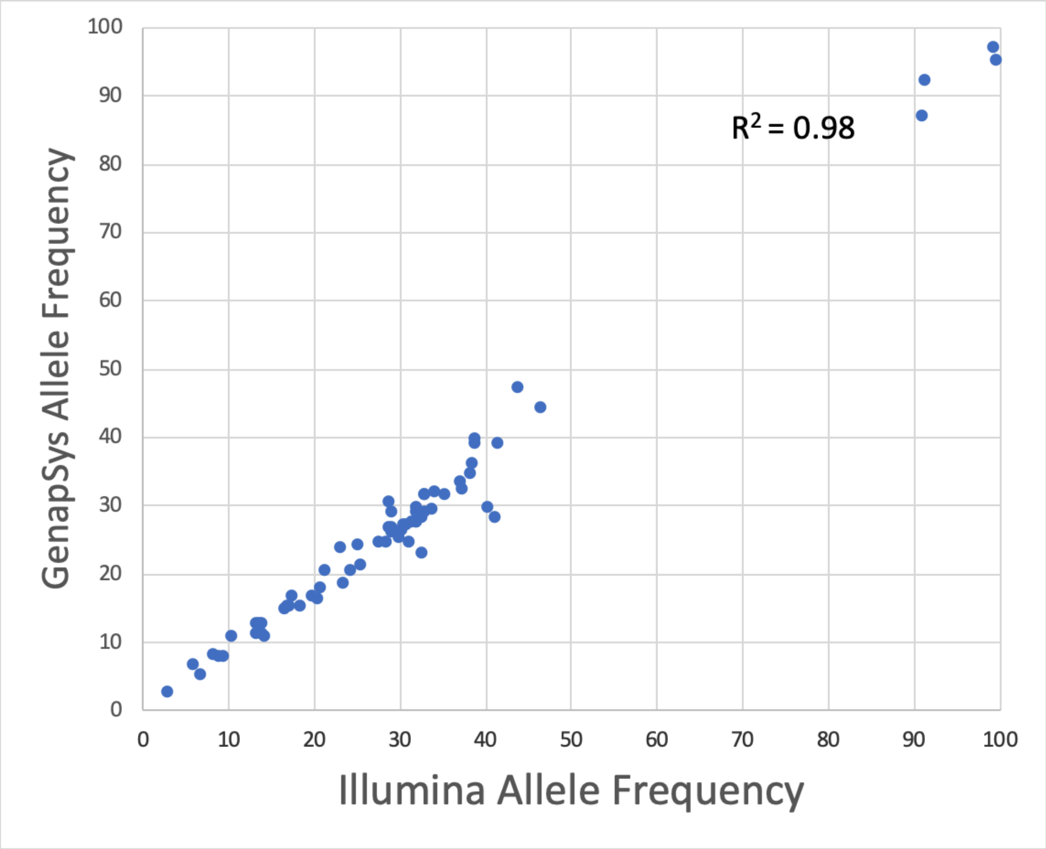

**SUPPLEMENTARY FIGURE 10:** Sensor Identification and Active Sensor Determination

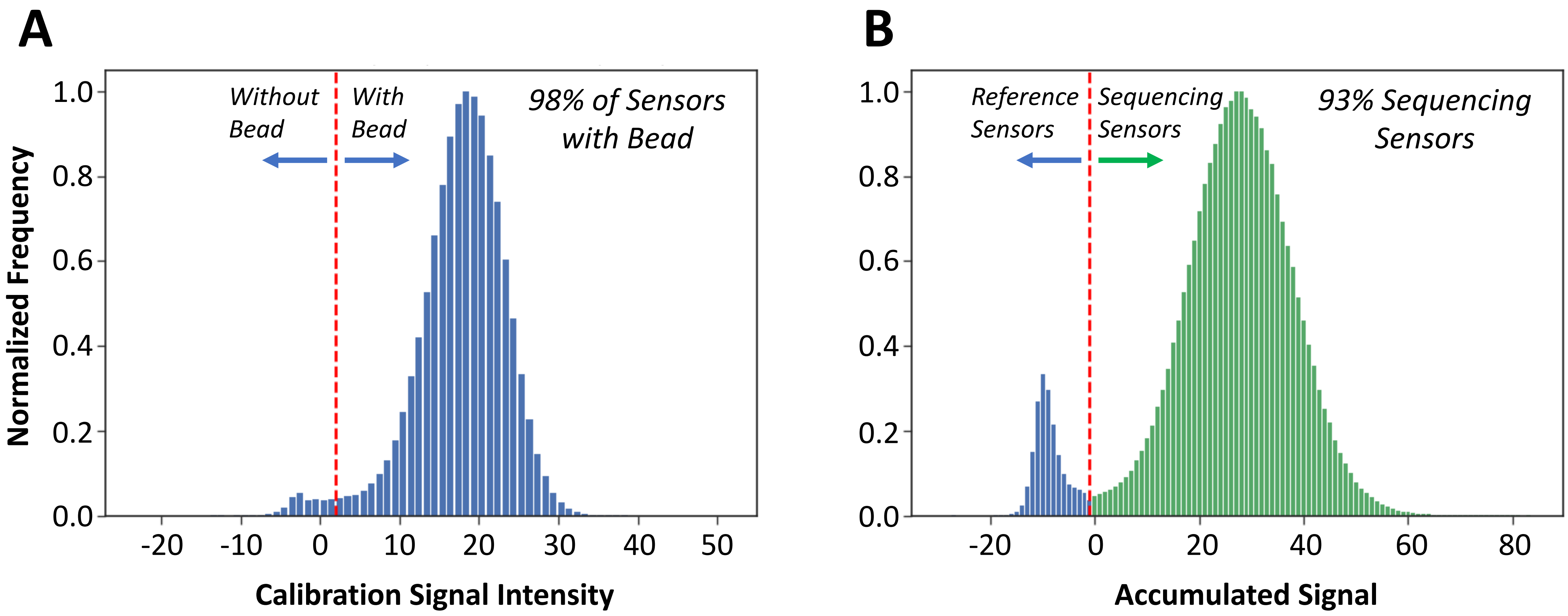

**SUPPLEMENTARY TABLE 1:** DNA Samples

| DNA Name | Description | Source | Catalog # | Genome Size | GC Content | Reference |
|---|---|---|---|---|---|---|
| *E. coli* | Escherichia coli K-12 | ATCC | 10798D-5 | 4.69 Mb | 51.5% | NARG00000000 |
| *C. jejuni* | Campylobacter jejuni NCTC11168 | ATCC | 700819D-5 | 1.64 Mb | 30.5% | AL111168 |
| *B. animalis* | Bifidobacterium animalis subsp. Lactis DSM10140 | DSM | 10140 | 1.94 Mb | 60.5% | CP001606 |
| NA12878 | Human CEPH/UTAH PEDIGREE 1463 | Coriell | NA12878 | 3.2 Gb | 40.8% | hg38 |
| HD701 | Quantitative Multiplex Reference Standard gDNA | Horizon Discovery | HD701 | 3.2 Gb | 40.8% | hg38 |
| HD827 | OncoSpan gDNA | Horizon Discovery | HD827 | 3.2 Gb | 40.8% | hg38 |

**SUPPLEMENTARY TABLE 2:** Targeted Sequencing

| | DNA Sample | | |
|---|---|---|---|
| **Statistic** | **NA12878** | **HD701** | **HD827** |
| Targeted Panel | xGen Exome Research (v1.0) | xGen Pan-Cancer (v1.5) | xGen Pan-Cancer (v1.5) |
| Panel Size | 39 Mb | 0.8 Mb | 0.8 Mb |
| On Target Rate | 80.5% | 64.4% | 79.2% |
| Average Coverage in Target Region | 115x | 624x | 1189x |
| Number of Reads | 55,645,073 | 11,542,744 | 10,593,385 |
| Number of Mapped Reads | 53,539,087 | 10,528,257 | 10,589,410 |
| Number of Mapped Reads to Targeted Region | 44,678,702 | 6,776,890 | 8,384,720 |
